# eQTL of KCNK2 regionally influences the brain sulcal widening: evidence from 15,597 UK Biobank participants with neuroimaging data

**DOI:** 10.1101/386821

**Authors:** Yann Le Guen, Cathy Philippe, Denis Riviere, Hervé Lemaitre, Antoine Grigis, Clara Fischer, Ghislaine Dehaene-Lambertz, Jean-François Mangin, Vincent Frouin

**Author notes:** Corresponding authors’ information: Address: CEA Saclay, Neurospin Bâtiment 145, 91191 Gif-sur-Yvette Cedex, France.

## Abstract

The grey and white matter volumes are known to reduce with age. This cortical shrinkage is visible on magnetic resonance images and is conveniently identified by the increased volume of cerebrospinal fluid in the sulci between two gyri. Here, we replicated this finding using the UK Biobank dataset and studied the genetic influence on these cortical features of aging. We divided all individuals genetically confirmed of British ancestry into two sub-cohorts (12,162 and 3,435 subjects for discovery and replication samples, respectively). We found that the heritability of the sulcal opening ranges from 15 to 45% (s.e.= 4.8%). We identified 4 new loci that contribute to this opening, including one that also affects the sulci grey matter thickness. We identified the most significant variant (rs864736) on this locus as being an expression quantitative trait locus (eQTL) for the KCNK2 gene. This gene regulates the immune-cell into the central nervous system (CNS) and controls the CNS inflammation, which is implicated in cortical atrophy and cognitive decline. These results expand our knowledge of the genetic contribution to cortical shrinking and promote further investigation into these variants and genes in pathological context such as Alzheimer’s disease in which brain shrinkage is a key biomarker.

## Introduction

The brain structure aspect alters throughout life. In particular, grey and white matter volumes are known to shrink with age in diseased and normal brains (Ge et al. 2002; Fjell and Walhovd 2010; Lockhart and DeCarli 2014). Numerous cross-sectional and longitudinal studies have confirmed this trend by either studying grey matter volume changes (Lemaitre et al. 2012) or cortical sulci widening (Kochunov et al. 2005; Shen et al. 2018). Importantly, the magnitude of brain shrinkage varies across regions and individuals, and increases with age (Raz et al. 2005). Multiple factors related to the environment or to genetics likely play a role in these changes. Such genetic effect is characterized in the hippocampus atrophy with the apolipoprotein E, ε4 allele (ApoE-ε4), which is also associated with an increased risk for developing late onset Alzheimer’s disease (AD) (Moffat et al. 2000). However, the genetic underpinnings of the brain sulcal features have not been investigated, except a few studies that were interested in the heritability of sulcal depth in extended pedigrees of young adults (Le Guen et al. 2017) or non-human primates (Kochunov et al. 2010). To the best of our knowledge, no genome wide association studies (GWAS) in imaging genetics with a sample size above 10,000 subjects were conducted on the brain sulcal features. In imaging-genetics, previous GWAS with such sample sizes have looked into the hippocampal and intracranial volumes (Stein et al. 2012), or the human subcortical brain structures (Hibar et al. 2015). These studies traditionally used meta-analyses which pooled subjects scanned in different centers with various scanners and from different age ranges. Such a meta-analysis initiative is best exemplified by the ENIGMA(Thompson et al. 2014) and CHARGE(Psaty et al. 2009) consortia. The UK Biobank project (Allen et al. 2012) offers a remarkable opportunity to address these issues by gathering data from a fairly homogenous population of subjects, and acquiring magnetic resonance images (MRI) on identical scanners operated at the same location. Additionally, it enables researchers to directly access a cohort with numerous participants while alleviating the uncertainty of meta-analyses.

In this paper, we consider ten prominent brain sulci that are automatically extracted and labelled using the Brainvisa cortical sulci recognition pipeline (Rivière et al. 2009; Perrot et al. 2011). These sulci are the central, the anterior and posterior cingulate, the inferior and superior temporal, the intraparietal, the subparietal, the superior and inferior frontal sulci and the Sylvian fissure (Mangin et al. 2015; Shen et al. 2018). Even though the subparietal sulcus is not as prominent as the others, it was included in our analysis because it lies in the precuneus which is a major target for atrophy in AD (Karas et al. 2007; Bailly et al. 2015). First, we replicated the known trends of cortical shrinking with age in a large sample of individuals from the UK Biobank, considering the grey matter thickness and sulcal opening. Second, we estimated the genetic influence on these features with the genome wide complex trait analysis (GCTA)(Yang et al. 2011). In this method the genetic relationship (kinship) matrix between subjects is computed to estimate the variance of an observed phenotype, which can be explained by the single nucleotide polymorphisms (SNPs), referred to as the heritability. Finally, we performed a genome wide association study (GWAS) of the phenotypes with the genotyped variants using PLINK (Purcell et al. 2007). A functional annotation of the phenotype-associated variants was then performed using the gene expression level published by the Genotype-Tissue Expression (GTEx) consortium (GTEx Consortium 2017), allowing the identification of expression quantitative trait loci (eQTLs).

## Methods

### Materials – Subjects

The present analyses were conducted under UK Biobank data application number 25251. The UK Biobank is a health research resource that aims to improve the prevention, diagnosis and treatment of a wide range of illnesses. Between the years 2006 and 2010, about 500,000 people aged between 45 and 73 years old, were recruited in the general population across Great Britain. In this work, we used the data released on January 2018, consisting of 20,060 subjects with a T1-weighted MRI. We included 15,040 in our discovery sample and 5,020 in our replication sample. The subjects were separated according to their availability in NIFTI format. In January 2018, replication sample was available only in DICOM format.

The UK Biobank genetic data underwent a stringent QC protocol, which was performed at the Wellcome Trust Centre for Human Genetics (Bycroft et al. 2017). We restrained our analysis to people identified by UK Biobank as belonging to the main white British ancestry subset (using the variable *in.white.British.ancestry.subset* in the file *ukb_sqc_v2.txt*). Additionally, we excluded from our analysis subjects with high missingness, high heterozygosity, first degree related individuals or sex mismatches. In total ~12,150 subjects in the discovery cohort and ~3,430 subjects in the replication cohort passed the image processing steps and the genetic criteria filtering. Both sets include approximately 48% of males and 52% of females.

### Cortical sulci extraction

The cortical sulci were extracted from T1-weighted images via the following steps. First, the brain mask was obtained with an automated skull stripping procedure based on the SPM8 skull-cleanup tool (Ashburner 2009). Second, the images were segmented into grey matter, white matter and cerebrospinal fluid using histogram scale-space analysis and mathematical morphology (Mangin et al. 2004). Third, individual sulci were segmented and labelled using Morphologist, the sulci identification pipeline from Brainvisa (version 4.5.1, (Rivière et al. 2009)). For segmentation, a kind of crevasse detector was used to reconstruct each fold geometry as the medial surface from the two opposing gyral banks that spanned from the most internal point of the fold to the convex hull of the cortex (Mangin et al. 2004). A Bayesian inspired pattern recognition approach relying on Statistical Probabilistic Anatomy Maps and multiscale spatial normalization was used to label the folds using a nomenclature of 125 sulci (Perrot et al. 2011; Mangin et al. 2015). For each sulcus, the average distance between both banks of the pial surface was used to quantify the sulcus width. This average distance was computed as the ratio between 1) the volume of CSF filling up the sulcus from the brain hull to the fold bottom and 2) the surface area of the sulcus estimated by half the sum of the areas of the triangles making up a mesh of the corresponding medial surface (**Figure 1a**). The average thickness of the cortical mantle on both sides of the sulcus was computed using a fast marching algorithm applied to a voxel-based binary representation of the cortex grey matter (Perrot et al. 2011).

**Figure 1.**
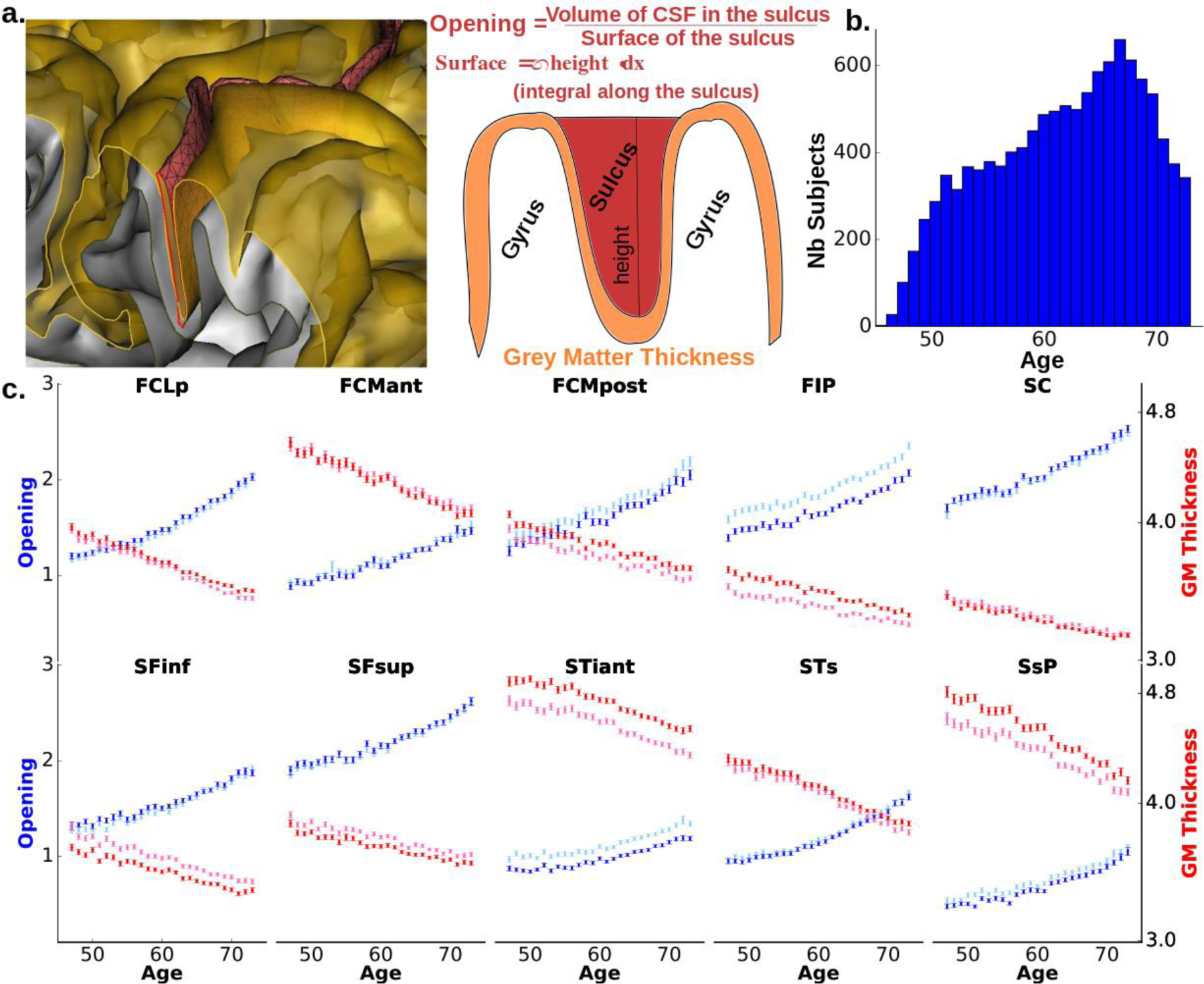
Evolution of sulcal opening and grey matter thickness with age in the main brain sulci. **a.** Schematic definition of the opening and grey matter thickness for the Brainvisa sulci. **b.** Age distribution in the UK Biobank sample with MR Imaging. **c.** Mean opening (red) and mean grey matter thickness (blue) vs age. (light blue and light red represent left hemisphere values; dark blue and red represent right hemisphere values)

### Age and sulci features relationship

In our discovery sample, to quantify the influence of age, we computed the mean of each sulcal feature for each age across the subjects with the same age. It is worth noting that the age at MRI scan is provided by UK Biobank with a one year precision. We excluded 45 and 46 year old subjects due to their small sample sizes. We can robustly estimate the mean feature per year after 50 years, because there are more than 200 subjects per 1-year interval. Last, using linear regression, we estimated the slope of a linear model adjusted between sulcal features and age (**Table S1**).

### Heritability estimation – Genome wide complex trait analyses (GCTA)

In our discovery sample, we used GCTA (Yang et al. 2011) that yields an estimate of the heritability (h²_SNPs_) in population studies with genotyped unrelated participants. We considered the genotyped SNPs variants common to the UKBiobank and UKBileve arrays (details at http://www.ukbiobank.ac.uk). In order to compute the kinship matrix of the population, specific SNPs were selected with PLINK v1.9(Purcell et al. 2007) using the following thresholds: missing genotype = 5% (70.783 variants excluded), Hardy-Weinberg equilibrium test (hwe) = 10^−6^ (11.318), and minor allele frequency (maf) = 1,0% (102.559). We kept the SNPs in moderate linkage disequilibrium with variation inflation factor 10 within a window of 50 SNPs (92.081 variants excluded). Then, we computed the genetic relationship matrix with GCTA using the 507.515 SNPs left. The amount of phenotypic variance captured by this matrix is estimated using a mixed-effects linear model. As covariates in our genetic analyses, we systematically included the gender, the genotyping array type, the age at the MRI session and the 10 genetic principal components provided by UK Biobank to account for population stratification. Correction for multiple comparisons were achieved using Bonferroni correction accounting for all our phenotypes and we retained as significant p < 0.00125 = 0.05/40 (2 hemispheres × 2 cortical features × 10 sulci). Using the GCTA Power Calculator (Visscher et al. 2014), the discovery cohort sample size provides above 99% statistical power to detect heritability values above 15%, with p < 0.00215.

### Genetic univariate association analyses

The genotype-phenotype association analyses were performed using PLINKv1.9 (Purcell et al. 2007), with the following thresholds: missing genotype = 10% (32.938 variants excluded), hwe = 10^−6^ (12.301), and maf = 1,0% (117.165), in total 621.852 variants passed the genetics QC for the discovery sample. Note that the missing genotype threshold is traditionally more stringent to estimate the kinship in GCTA analysis than to perform GWAS. The genome wide significant threshold for the discovery sample was set to p < 5·10^−8^ and p < 0.05 in the replication sample for variants that passed the first threshold.

## Results

### Age-related cortical shrinking

The UK Biobank large sample size enabled us to precisely estimate the mean of sulcal opening and grey matter (GM) thickness per age with a 1-year precision. **Figure 1** (**c, d**) underlines a strong correlation between the age and these two cortical features in the discovery sample. Between 47 and 73 years old, the sulcal opening increases on average of 0.025 mm/year, while the GM thickness decreases on average of 0.015 mm/year (**Table S1**). The Sylvian fissure and the subparietal sulcus show the maximum increase of opening and decrease of GM thickness, respectively, while the inferior temporal and left intraparietal have the minimum.

### Heritability of sulcal opening and grey matter thickness around the sulci

In the discovery sample, we estimated the heritability (**h²_SNPs_**) of the sulcal opening and GM thickness using the GCTA method (Yang et al. 2011), i.e. the variance explained by all the SNPs (**Figure 2**). Significant **h²_SNPs_** estimates for the sulcal opening range from 0.15 to 0.45, with a minimum in the left inferior temporal and a maximum in the left central sulcus (**Table S2**). Significant **h²_SNPs_** estimates for the GM thickness range from 0.15 to 0.37, with a minimum in the left superior frontal and a maximum in the left Sylvian fissure. Note that the sulcal opening heritability values are all higher than the ones of the GM thickness.

**Figure 2.**
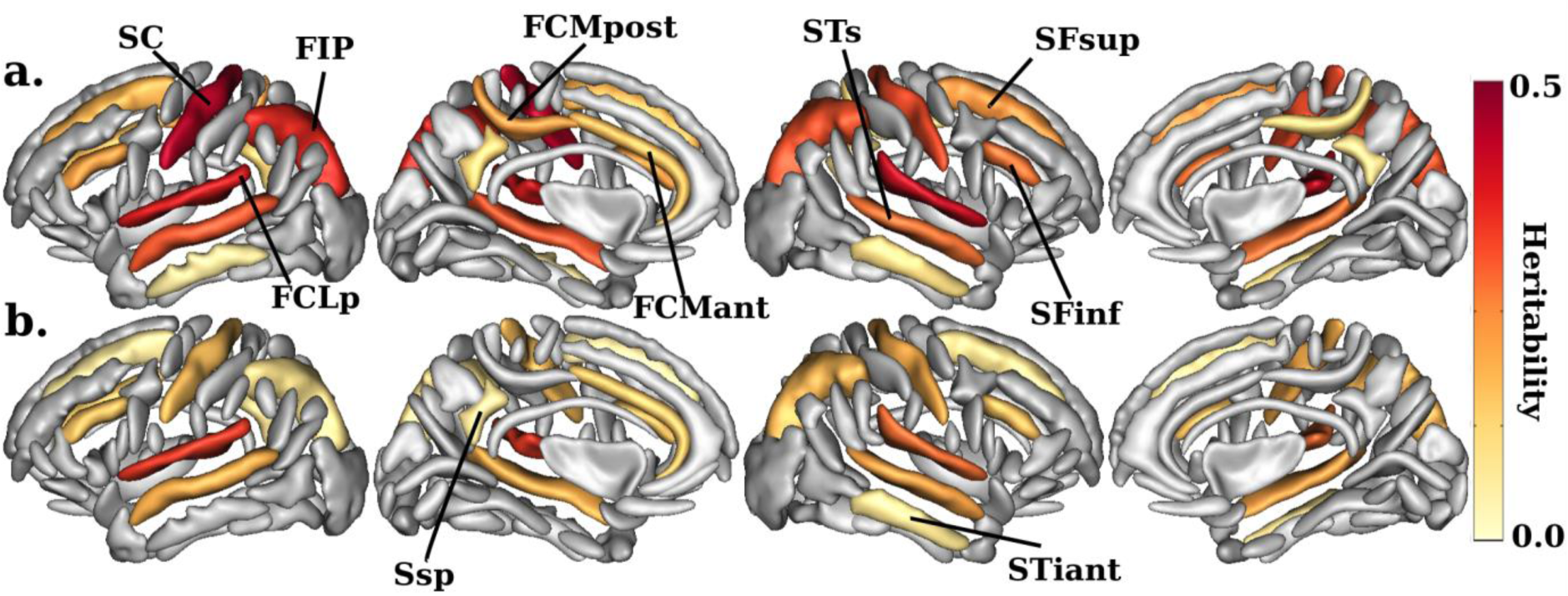
Heritability estimates of the sulcal opening (a.) and sulcal grey matter thickness (b.). (p < 0.00125 Bonferroni corrected). Main sulci have been annotated using Brainvisa abbreviation (**SC**: Central, **FIP**: Intraparietal, **FCLp**: Sylvian fissure, **FCMpost**: Posterior Cingulate, **FCMant**: Anterior Cingulate, **STs**: Superior Temporal, **STiant**: Inferior Temporal, **SFsup**: Superior Frontal, **SFinf**: Inferior Frontal, **Ssp**: Subparietal). The sulci are displayed using the Statistical Probability Anatomy Map (SPAM) representation, which represents the average sulci shape and position on the reference base of the Brainvisa sulci extraction pipeline (Perrot et al. 2011).

### Genome-wide association study (GWAS) of the cortical features

We performed a genome-wide association study on the genotyped data for the sulcal opening and GM thickness in the ten sulci. Manhattan and QQ plots are shown in the supplementary materials (**Figures S1-4**). **Table 1** summarizes the 24 phenotype-SNP associations that were genome-wide significant in the discovery sample and nominally significant in the replication sample. Among these associations, 12 SNPs were unique at 5 different loci. The most represented locus, with 17 replicated association hits, is on chromosome 1, 27kb before the transcription start site of the KCNK2. Within this locus, the two main associated SNPs are rs59084003 and rs864736 with 7 and 4 significant hits respectively. It should be noted that these two SNPs are not in strong linkage disequilibrium (LD) and other significant SNPs on this locus are either in LD with the first or the second one (**Figure S5**). The second most represented locus, with 4 significant associations, is on chromosome 16. This locus lies in the starting region of the non-coding RNA LOC101928708, in the vicinity of the protein coding gene C16orf95, followed by FBXO31. On this locus, the main associated SNP is rs9933149 with 3 hits. The three other loci each include a single significant replicated phenotype-SNP association. On chromosome 8, rs11774568, associated with the GM thickness in the left Sylvian fissure, is in a region with a high density of genes, between the genes DEFB136 and DEFB135. On chromosome 9, rs10980645, associated with the central sulcus opening, is an intronic variant of the LPAR1 gene. On chromosome 12, rs12146713, associated with the right STs opening, is an intronic variant of the NUAK1 gene.

**Table 1.**
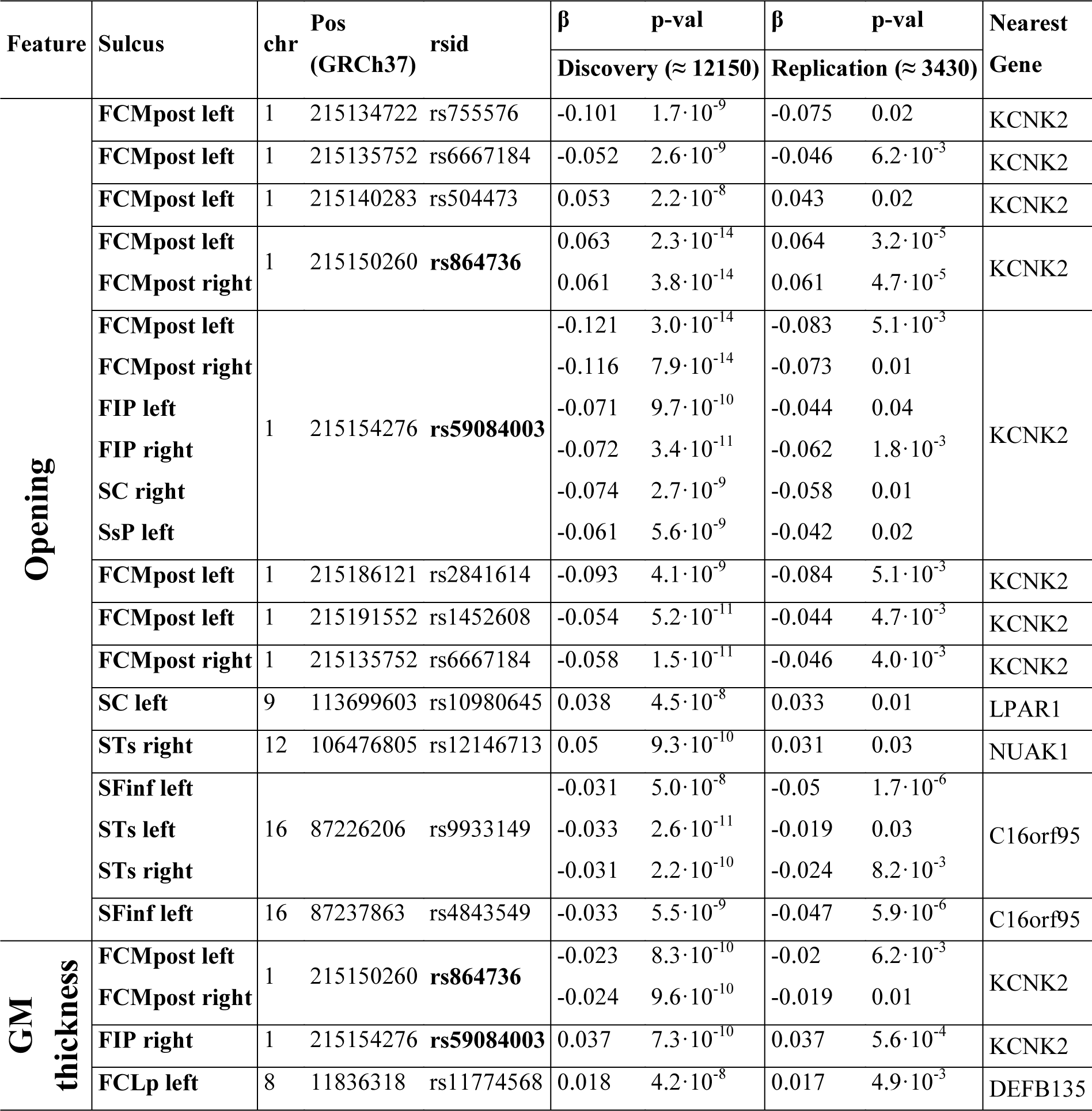
Significant genome wide association hit SNPs. (discovery p < 5·10^−8^ and replication p < 0.05). Bold rsid correspond to variants that are further investigated in **Figure 3**, the remainders are in **Figures S7-S8**.

### Regional significance and direction of effect of the hit variants

DNA region upward KCNK2 harbors significant SNP association with the sulcal opening (**Figure 3**) and GM thickness (**Figure S6**) of the posterior cingulate, the central and intraparietal sulci. In addition, the implicated SNPs show near genome-wide significant influence on the sulcal features in the superior temporal, inferior frontal and subparietal sulci. Overall, this suggests a brain wide regulation for this genomic region. The sulcal opening is increased in carriers of the minor allele of rs864736 (maf = 46%, in our discovery sample), while it is decreased in carriers of the minor allele of rs59084003 (maf = 7%). The locus on chromosome 16 displays a more specific spatial control over the temporal and frontal lobes, with significant sulci including the inferior frontal, the superior and inferior temporal and the Sylvian fissure (**Figures S7-8 a.**). The sucal opening of these sulci is decreased in carriers of the minor allele rs9933149 (maf = 38%). It should be noted that the SNPs of these two loci have a significant pleiotropic influence on sulcal opening and GM thickness.

**Figure 3.**
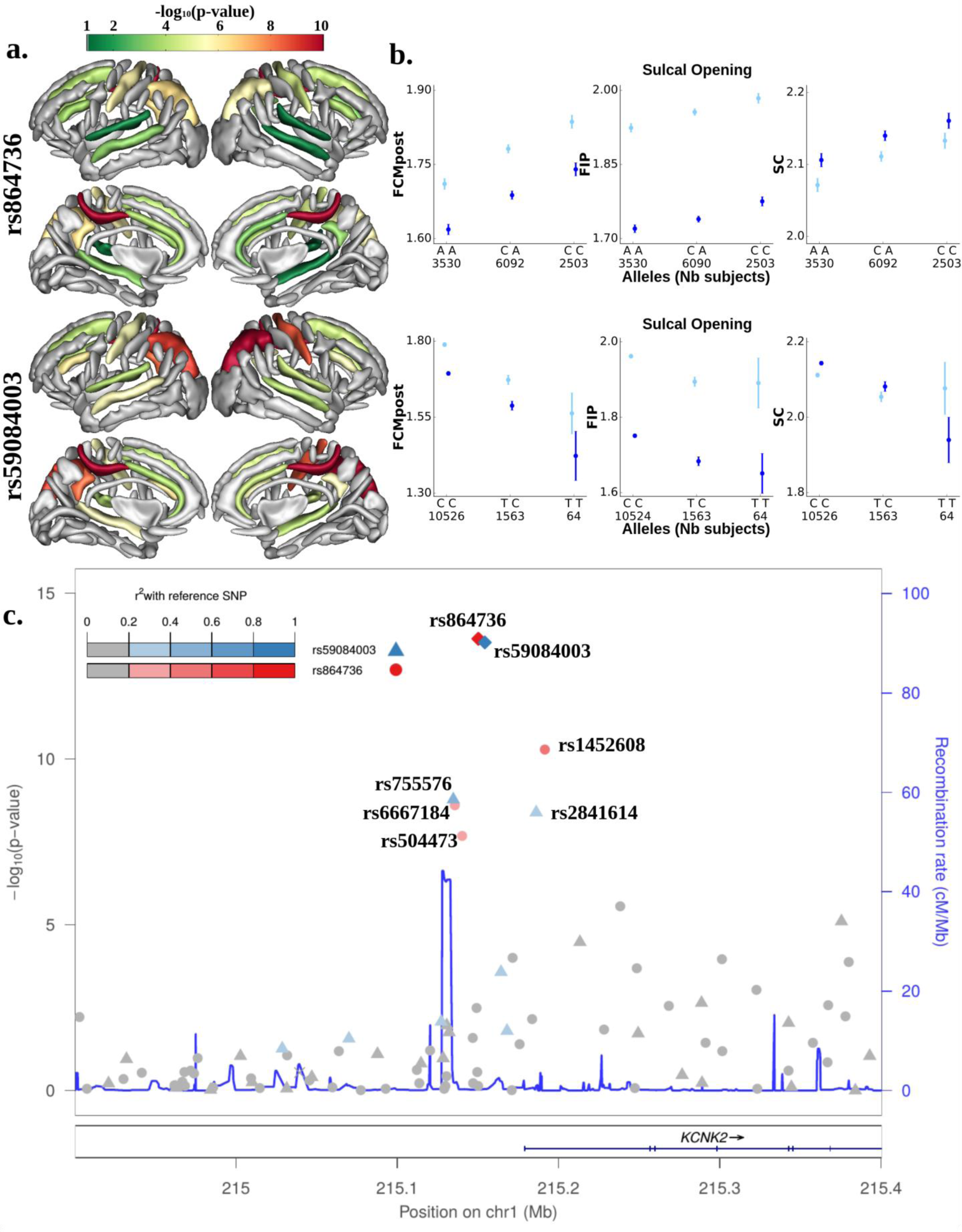
GWAS hits upstream of KCNK2 regulating the sulcal opening. First and second lines correspond to rs864736 and rs59084003, respectively. Lines represents respectively: **a)** the log10(p-value) of each SNPs mapped onto the nominally significant sulci among the ten considered; **b)** the mean sulcal opening and standard error for each configuration of variants in the most significant sulci; **c)** Locuszoom display (Pruim et al. 2011) of the phenotype-variants association for the region upstream of KCNK2 with the left posterior cingulate sulcus opening as a phenotype.

Regarding the three remaining loci, they preferentially influence one out of the two sulcal features. The intronic variant (rs12146713) of NUAK1 significantly influences the sulcal opening in the temporal region (Sylvian fissure, superior and inferior temporal sulci), as well as in the inferior frontal and intraparietal sulci (**Figures S7-8 b.**). The intronic variant (rs10980645) of LPAR1 significantly affects the sulcal opening of the central sulcus, and moderately affects the posterior cingulate (**Figures S7-8 c.**). The variant (rs11774568) chromosome 8, near DEFB136, appears to be linked with the GM thickness of the Sylvian fissure and superior temporal sulcus (**Figures S7-8 d.**).

### Functional annotation of the loci

To obtain information on the role of the region upstream of KCNK2, we investigated the gene expression QTL using the GTEx database (GTEx Analysis Release V7 (dbGaP Accession phs000424.v7.p2)) (GTEx Consortium 2017). We found that the variants rs864736 and rs14526008, which are in LD, are significant eQTLs for the KCNK2 gene expression, with significant multi-tissue meta-analysis RE2 (random-effects model 2) (Han and Eskin 2011) p-values of 9.7·10^−6^ and 5.7·10^−9^, respectively. **Figure 4** presents the effect size in various tissues of rs864736 allelic configuration on KCNK2 gene expression. Even though the association barely reaches nominal significance in single brain tissue due to low sample sizes (~80-140 subjects), it overall emphasizes strong effects across brain tissues. The other significantly associated SNP near KCNK2 (rs59084003) was not reported as a significant eQTL in GTEx possibly for technical reasons because of its small minor allele frequency (7%). Indeed, the effect of allelic configurations cannot be well observed in the relative small GTEx sample. **Table S3** summarizes the other eQTLs found in GTEx for the different loci.

**Figure 4.**
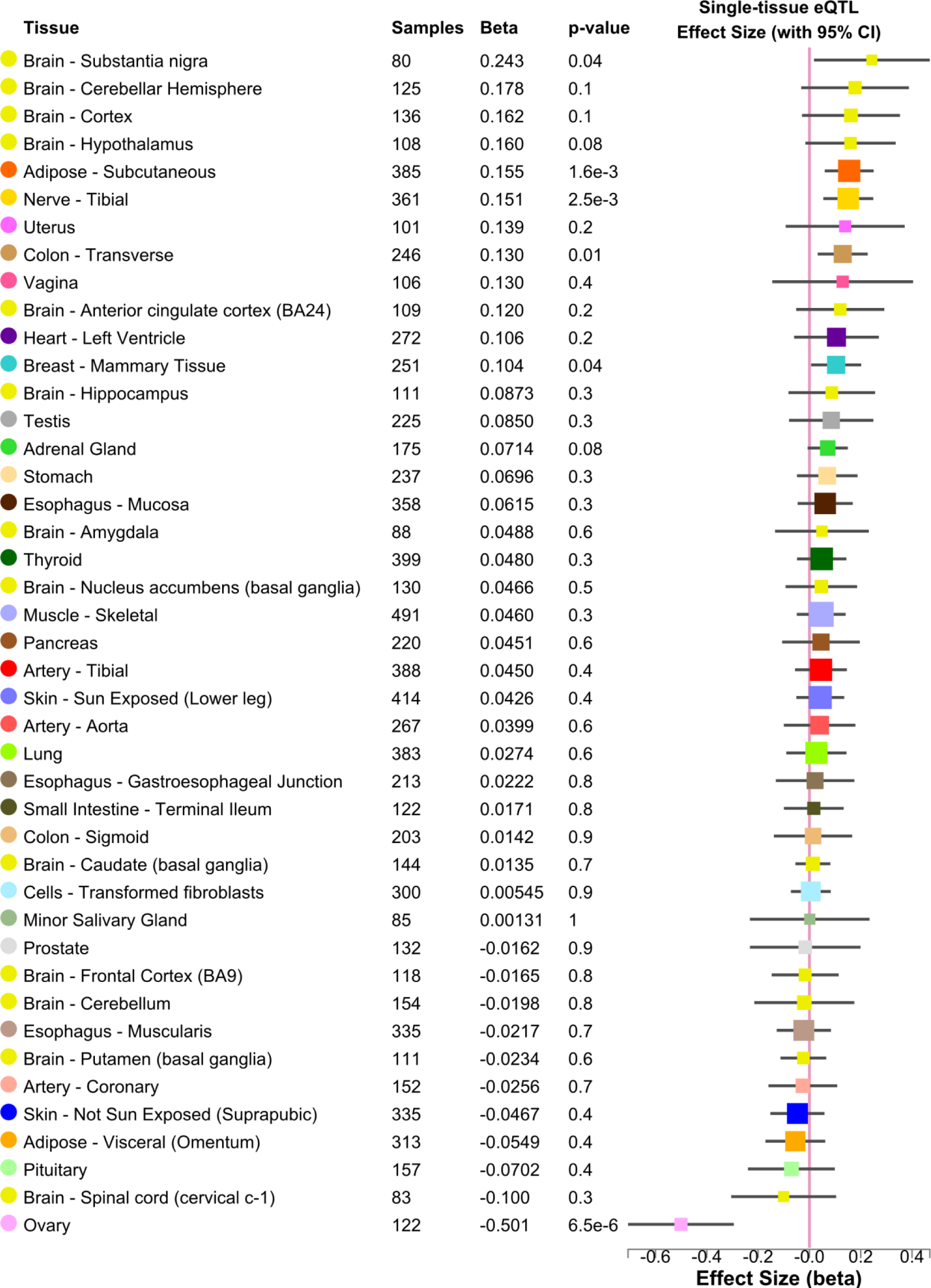
Multi-tissue eQTLs comparison for gene KCNK2 and variant rs864736. (ENSG00000082482.9 KCNK2 and 1_215150260_C_A_b37 eQTL). Meta-Analysis Random Effect Model2 (Han and Eskin 2011) p-val = 9.7·10^−6^. (Data Source: GTEx Analysis Release V7, dbGaP Accession phs000424.v7.p2)

## Discussion

The bulk of participants in the UK Biobank are older than 50 years old and results have to be discussed within this interval. In order to scrutinize certain genetic effects on brain features, it is important to understand the effect of age. Indeed, some genetic influences may only be revealed in a relatively aged cohort such as the UK Biobank. In this study, we emphasized a steady age effect on the sulcal opening and GM thickness between 45 and 75 years of age. Furthermore, we showed that these brain features are genetically controlled by estimating their heritability in ten prominent sulci and we also identified several of their causal variants.

It is well known that with age, the GM and white matter (WM) volumes decreases (Raz et al. 2005), while the amount of cerebrospinal fluid (CSF) in the cortical folds increases (Good et al. 2001). In our analysis, we confirmed the well-established results regarding the GM thickness decrease and sulcal opening increase with age in the UK Biobank cohort. The assessment of cortical sulci opening is well described and correlates with neurocognitive decline in mild cognitive impairment and dementia disorders (Bastos Leite et al. 2004). The cortical sulci widening with age is likely related to the reduction of gyral thickness resulting in the dilatation of the sulci (Magnotta 1999; Symonds et al. 1999), but could also account for neurodegenerative processes occurring in the underlying white matter (Gunning-Dixon et al. 2009). While, the GM thickness and sulcal widening are correlated (Kochunov et al. 2008; Liu et al. 2013), the robustness of the two measures might differ. We obtained less significant variant-phenotype associations in GM thickness, because the sulcal opening might be more consistently measured across individuals than the GM thickness. This might be due to the fact that the MRI contrast between GM and CSF remains more stable across the lifespan than the GM / WM contrast (Kochunov et al. 2005). Thus, sulcal widening is commonly used by radiologists as a surrogate of cortical atrophy in clinical settings (Shen et al. 2018). It could also reflect a higher sensitivity to the primal effects of aging because of the consequences of grey and white matter atrophy. Note that the opening of larger sulci like the Sylvian valley could also be impacted by global mechanical compensation for aging processes.

The main finding of this paper is that the locus upstream of the KCNK2 transcription start site influences the sulcal opening and GM thickness. Additionally, the tissue specific gene expression (eQTL) analysis of the GTEx consortium emphasizes that overall (meta-analysis of all tissues), and particulary in the brain, this DNA region regulates the expression of KCNK2 (GTEx Consortium 2017). Thus, we can legitimately assume a link between the regulation of KCNK2 expression and the amplitude of sulcal opening. In other words, depending on the allelic configurations in the region upstream of KCNK2, an individual will have his sulci comparatively enlarged. Because sulcal widening is a marker of cortical atrophy as we pinpointed in the previous paragraph, there is a potential link between KCNK2 expression level and brain atrophy. The KCNK2 gene, also known as TREK1, is a member of the two-pore-domain potassium channel family which is expressed predominantly in the brain (Hervieu et al. 2001). Previous literature emphasized several functions for KCNK2 gene in the brain. First, the KCNK2 regulates the blood-brain barrier function and inflammation in the brain of mice (Bittner et al. 2013) and humans (Bittner et al. 2014). The inhibition or deletion of KCNK2 facilitates lymphocytes migration into the central nervous system (CNS) and promotes autoimmune CNS inflammation (Bittner et al. 2013). Second, in mice, the knockdown of KCNK2 gene impairs the neuronal migration of late-born cortical excitatory neurons, which are precursors of Layer II/III neurons (Bando et al. 2014). Third, in rat hippocampal astrocytes, the increase of KCNK2 expression mediates neuroprotection during ischemia (Banerjee et al. 2016). The mechanism might involve KCNK2 blockade, inhibiting neuronal apoptosis and protecting the brain from cerebral ischemic injury (Wang et al. 2018). Finally, KCNK2 over expression was shown to exacerbate memory impairment in middle-age mice (Cai et al. 2017). To summarize, KCNK2 controls several major cellular responses involved in memory formation and is believed to participate in neuroinflammation, cerebral ischemia and blood-brain barrier dysfunction (Bittner et al. 2014; Cai et al. 2017; Wang et al. 2018).

The first role suggests the most promising direction of future work, because previous studies have proposed that neuroinflammation is involved in cognitive decline in midlife (Marsland et al. 2015) and implicated in pathological age-related changes and AD (McGeer and McGeer 2001). Throughout life, stress, recurrent inflammation and subclinical cerebrovascular events potentially contribute to brain aging (Raz and Rodrigue 2006). The link between our findings and inflammation indicates a potential mediation role for KCNK2. Finally, it is difficult to disentangle whether or not brain inflammation has a deleterious role on cognitive functions, since there is no clear consensus. A recent study however emphasizes a slower progression of AD in patients with early neuroinflammation (Hamelin et al. 2016).

In conclusion, in a sample of 15,597 subjects representative of the general population of British ancestry, we have shown that an eQTL of KCNK2 influences sulcal widening. This appears coherent with the role of KCNK2, which affects the regulation of inflammation response in the brain.

## ETHICAL STATEMENT

### Compliance with Ethical Standards

The authors declare that they comply with the Ethical Standards of the journal

### Informed consent

Informed consent was obtained from all individual participants included in the study.

### Ethical approval

All procedures performed in studies involving human participants were in accordance with the ethical standards of the institutional and/or national research committee and with the 1964 Helsinki declaration and its later amendments or comparable ethical standards.

## References

Allen N, Sudlow C, Downey P, et al (2012) UK Biobank: Current status and what it means for epidemiology. Heal Policy Technol 1:123–126. doi: 10.1016/j.hlpt.2012.07.003

Ashburner J (2009) Computational anatomy with the SPM software. Magn Reson Imaging 27:1163–1174. doi: 10.1016/j.mri.2009.01.006

Bailly M, Destrieux C, Hommet C, et al (2015) Precuneus and Cingulate Cortex Atrophy and Hypometabolism in Patients with Alzheimer&#x2019;s Disease and Mild Cognitive Impairment: MRI and 18F-FDG PET Quantitative Analysis Using FreeSurfer, Precuneus and Cingulate Cortex Atrophy and Hypometabolism in P. BioMed Res Int BioMed Res Int 2015, 2015:e583931. doi: 10.1155/2015/583931, 10.1155/2015/583931

Bando Y, Hirano T, Tagawa Y (2014) Dysfunction of KCNK potassium channels impairs neuronal migration in the developing mouse cerebral cortex. Cereb Cortex 24:1017– 1029. doi: 10.1093/cercor/bhs387

Banerjee A, Ghatak S, Sikdar SK (2016) l-Lactate mediates neuroprotection against ischaemia by increasing TREK1 channel expression in rat hippocampal astrocytes in vitro. J Neurochem 265–281. doi: 10.1111/jnc.13638

Bastos Leite AJ, Scheltens P, Barkhof F (2004) Pathological Aging of the Brain. Top Magn Reson Imaging 15:369–389. doi: 10.1097/01.rmr.0000168070.90113.dc

Bittner S, Ruck T, Fernández-Orth J, Meuth SG (2014) TREK-king the blood-brain-barrier. J Neuroimmune Pharmacol 9:293–301. doi: 10.1007/s11481-014-9530-8

Bittner S, Ruck T, Schuhmann MK, et al (2013) Endothelial TWIK-related potassium channel-1 (TREK1) regulates immune-cell trafficking into the CNS. Nat Med 19:1161– 1165. doi: 10.1038/nm.3303

Bycroft C, Freeman C, Petkova D, et al (2017) Genome-wide genetic data on ~500,000 UK Biobank participants. bioRxiv 166298. doi: 10.1101/166298

Cai Y, Peng Z, Guo H, et al (2017) TREK-1 pathway mediates isoflurane-induced memory impairment in middle-aged mice. Neurobiol Learn Mem 145:199–204. doi: 10.1016/j.nlm.2017.10.012

Fjell AM, Walhovd KB (2010) Structural Brain Changes in Aging: Courses, Causes and Cognitive Consequences. Rev Neurosci 21:187–221. doi: 10.1515/REVNEURO.2010.21.3.187

Ge Y, Grossman RI, Babb JS, et al (2002) Age-related total gray matter and white matter changes in normal adult brain. Part I: volumetric MR imaging analysis. Am J Neuroradiol 23:1327–33

Good CD, Johnsrude I, Ashburner J, et al (2001) Cerebral asymmetry and the effects of sex and handedness on brain structure: A voxel-based morphometric analysis of 465 normal adult human brains. Neuroimage 14:685–700. doi: 10.1006/nimg.2001.0857

GTEx Consortium (2017) Genetic effects on gene expression across human tissues. Nature 550:2041–213. doi: 10.1038/nature24277

Gunning-Dixon FM, Brickman AM, Cheng JC, Alexopoulos GS (2009) Aging of cerebral white matter: a review of MRI findings. Int J Geriatr Psychiatry 24:109–117. doi: 10.1002/gps.2087

Hamelin L, Lagarde J, Dorothée G, et al (2016) Early and protective microglial activation in Alzheimer’s disease: a prospective study using 18 F-DPA-714 PET imaging. Brain 139:1252–1264. doi: 10.1093/brain/aww017

Han B, Eskin E (2011) Random-effects model aimed at discovering associations in meta-analysis of genome-wide association studies. Am J Hum Genet 88:586–598. doi: 10.1016/j.ajhg.2011.04.014

Hervieu GJ, Cluderay JE, Gray CW, et al (2001) Distribution and expression of TREK-1, a two-pore-domain potassium channel, in the adult rat CNS. Neuroscience 103:899–919. doi: 10.1016/S0306-4522(01)00030-6

Hibar DP, Stein JL, Renteria ME, et al (2015) Common genetic variants influence human subcortical brain structures. Nature 520:224–229. doi: 10.1038/nature14101

Karas G, Scheltens P, Rombouts S, et al (2007) Precuneus atrophy in early-onset Alzheimer’s disease: A morphometric structural MRI study. Neuroradiology 49:967–976. doi: 10.1007/s00234-007-0269-2

Kochunov P, Glahn DC, Fox PT, et al (2010) Genetics of primary cerebral gyrification: Heritability of length, depth and area of primary sulci in an extended pedigree of Papio baboons. Neuroimage 53:1126–1134. doi: 10.1016/j.neuroimage.2009.12.045

Kochunov P, Mangin JF, Coyle T, et al (2005) Age-related morphology trends of cortical sulci. Hum Brain Mapp 26:210–220. doi: 10.1002/hbm.20198

Kochunov P, Thompson PM, Coyle TR, et al (2008) Relationship among neuroimaging indices of cerebral health during normal aging. Hum Brain Mapp 29:36–45. doi: 10.1002/hbm.20369

Le Guen Y, Auzias G, Leroy F, et al (2017) Genetic Influence on the Sulcal Pits: On the Origin of the First Cortical Folds. Cereb Cortex 1–12. doi: 10.1093/cercor/bhx098

Lemaitre H, Goldman AL, Sambataro F, et al (2012) Normal age-related brain morphometric changes: nonuniformity across cortical thickness, surface area and gray matter volume? Neurobiol Aging 33:617.e1–617.e9. doi: 10.1016/j.neurobiolaging.2010.07.013

Liu T, Sachdev PS, Lipnicki DM, et al (2013) Limited relationships between two-year changes in sulcal morphology and other common neuroimaging indices in the elderly. Neuroimage 83:12–17. doi: 10.1016/j.neuroimage.2013.06.058

Lockhart SN, DeCarli C (2014) Structural Imaging Measures of Brain Aging. Neuropsychol Rev 24:271–289. doi: 10.1007/s11065-014-9268-3

Magnotta V a (1999) Quantitative In Vivo Measurement of Gyrification in the Human Brain: Changes Associated with Aging. Cereb Cortex 9:151–160. doi: 10.1093/cercor/9.2.151

Mangin J-F, Perrot M, Operto G, et al (2015) Sulcus Identification and Labeling. Elsevier Inc.

Mangin JF, Rivière D, Cachia A, et al (2004) A framework to study the cortical folding patterns. Neuroimage 23:129–138. doi: 10.1016/j.neuroimage.2004.07.019

Marsland AL, Gianaros PJ, Kuan DCH, et al (2015) Brain morphology links systemic inflammation to cognitive function in midlife adults. Brain Behav Immun 48:195–204. doi: 10.1016/j.bbi.2015.03.015

McGeer PL, McGeer EG (2001) Inflammation, autotoxicity and Alzheimer disease. Neurobiol Aging 22:799–809. doi: 10.1016/S0197-4580(01)00289-5

Moffat SD, Szekely CA, Zonderman AB, et al (2000) Longitudinal change in hippocampal volume as a function of apolipoprotein E genotype. Neurology 55:134–136. doi: 10.1212/WNL.55.1.134

Perrot M, Rivière D, Mangin J-F (2011) Cortical sulci recognition and spatial normalization. Med Image Anal 15:529–50. doi: 10.1016/j.media.2011.02.008

Pruim RJ, Welch RP, Sanna S, et al (2011) LocusZoom: Regional visualization of genome-wide association scan results. Bioinformatics 27:2336–2337. doi: 10.1093/bioinformatics/btq419

Psaty BM, O’Donnell CJ, Gudnason V, et al (2009) Cohorts for Heart and Aging Research in Genomic Epidemiology (CHARGE) Consortium design of prospective meta-analyses of genome-wide association studies from 5 Cohorts. Circ Cardiovasc Genet 2:73–80. doi: 10.1161/CIRCGENETICS.108.829747

Purcell S, Neale B, Todd-Brown K, et al (2007) PLINK: A Tool Set for Whole-Genome Association and Population-Based Linkage Analyses. Am J Hum Genet 81:559–575. doi: 10.1086/519795

Raz N, Lindenberger U, Rodrigue KM, et al (2005) Regional brain changes in aging healthy adults: General trends, individual differences and modifiers. Cereb Cortex 15:1676– 1689. doi: 10.1093/cercor/bhi044

Raz N, Rodrigue KM (2006) Differential aging of the brain: Patterns, cognitive correlates and modifiers. Neurosci Biobehav Rev 30:730–748. doi: 10.1016/j.neubiorev.2006.07.001

Rivière D, Geffroy D, Denghien I, et al (2009) BrainVISA: an extensible software environment for sharing multimodal neuroimaging data and processing tools. Neuroimage 47:S163. doi: 10.1016/S1053-8119(09)71720-3

Shen X, Liu T, Tao D, et al (2018) Variation in longitudinal trajectories of cortical sulci in normal elderly. Neuroimage 166:1–9. doi: 10.1016/j.neuroimage.2017.10.010

Stein JL, Medland SE, Vasquez AA, et al (2012) Identification of common variants associated with human hippocampal and intracranial volumes. Nat Genet 44:552–61. doi: 10.1038/ng.2250

Symonds LL, Archibald SL, Grant I, et al (1999) Does an Increase in Sulcal or Ventricular Fluid Predict Where Brain Tissue Is Lost? J Neuroimaging 9:201–209. doi: 10.1111/jon199994201

Thompson PM, Stein JL, Medland SE, et al (2014) The ENIGMA Consortium: Large-scale collaborative analyses of neuroimaging and genetic data. Brain Imaging Behav 8:153– 182. doi: 10.1007/s11682-013-9269-5

Visscher PM, Hemani G, Vinkhuyzen A a E, et al (2014) Statistical power to detect genetic (co) variance of complex traits using SNP data in unrelated samples. PLoS Genet 10:e1004269. doi: 10.1371/journal.pgen.1004269

Wang W, Liu D, Xiao Q, et al (2018) Lig4-4 selectively inhibits TREK-1 and plays potent neuroprotective roles in vitro and in rat MCAO model. Neurosci Lett 671:93–98. doi: 10.1016/j.neulet.2018.02.015

Yang J, Lee SH, Goddard ME, Visscher PM (2011) GCTA: A Tool for Genome-wide Complex Trait Analysis. Am J Hum Genet 88:76–82. doi: 10.1016/j.ajhg.2010.11.011

